# CRITICAL ROLE OF ADULT-BORN DENTATE GRANULE NEURONS IN PATTERN COMPLETION’

**DOI:** 10.1101/2025.05.15.654267

**Authors:** P Roullet, M Koehl, P Delage, F Guillemot, DN Abrous

## Abstract

The dentate gyrus contains neurons generated during development and adulthood, yet their distinct roles remain unclear. Using a transgenic mouse model, we show that adult-born neurons contribute to pattern completion, essential for episodic memory. This has implications for aging, PTSD, and addiction, where excessive pattern completion leads to maladaptive memories. Targeting ABNs could offer new therapeutic strategies to regulate memory processes and improve mental health.

---

The recollection of memories relies on several interconnected cognitive processes. After encoding—during which new information is integrated with existing knowledge—memories are stabilized and stored over time. Accessing and reconstructing past experiences depends on various mechanisms, including pattern completion, a process that allows the brain to retrieve a complete memory from partial or incomplete cues (Rolls, 1996;Rolls, 2013).

The dentate gyrus, particularly neurons generated during adulthood, has been implicated in memory recollection. Research has shown that ablating, silencing or inhibiting these neurons impairs memory retrieval (Arruda-Carvalho *et al*., 2011;Lods *et al*., 2021;Masachs *et al*., 2021), while stimulating their activity during retrieval enhances the accuracy of long-term memory recall (Lods *et al*., 2022). Although these findings suggest that adult-born neurons are both necessary and sufficient for accessing intact stored memories, their role in reconstructing a complete memory from partial cues has not been yet investigated.

To fill this gap, we used an already established inducible transgenic mouse model to stop the production of new neurons (Andersen *et al*., 2014;Castro and Guillemot, 2011). In this model, Ascl1, a bHLH transcription factor involved in cell cycle exit and proliferation, is deleted in radial glial cells by crossing mice carrying the transgenes *Glast-CreERT2* and *Ascl1* ^*fl/fl*^ and administering tamoxifen (Tam) to the double transgenic mice. Adult neurogenesis was examined in *Glast-CreERT2*^*-/-*^, *Ascl1* ^*+/+*^ (WT); *Glast-CreERT*^*+/-*^, *Ascl1* ^*+/+*^ (Cre+); *Glast-CreERT2*^*-/-*^, *Ascl1* ^*f/f*^ (FF); and *Glast-CreERT2*^*+/-*^, *Ascl1* ^*fl/fl*^ (cKO) mice treated with Tam at 2 months of age. CldU was injected 1 month later to verify the efficacy of Tam treatment at this early time point. Three months after CldU injection, we analyzed the number of CldU label-retaining cells and the number of immature neurons expressing doublecortin (DCX).

As expected, the number of CldU-label-retaining cells was dramatically decreased in cKO mice compared to the other groups. This confirms that one month after Tam injections, neurogenesis was abolished in cKO mice (Figure 1A, [Cre effect F_1,24_=0.045, p=ns; Flox effect F_1,24_=38.14, p<0.0001, Cre x Flox interaction F_1,24_=4.632, p<0.05 with WT=Cre+ > cKO at p<0.001]). In FF mice, which do not carry the *Glast-CreERT2* construct, the number of CldU-IR cells was lower than in WT and Cre+ groups [p=0.01 an p=0.0003, respectively], although it did not significantly differ from the cKO mice (p=0.11). This phenomenon previously observed (Andersen *et al*., 2014) suggests that the *Ascl1* ^*fl/fl*^ allele is hypomorphic and expresses *Ascl1* at a level lower than WT, leading to reduced neurogenesis in *Ascl1* ^*fl/fl*^ animals even in the absence of recombination.

**Figure 1.**
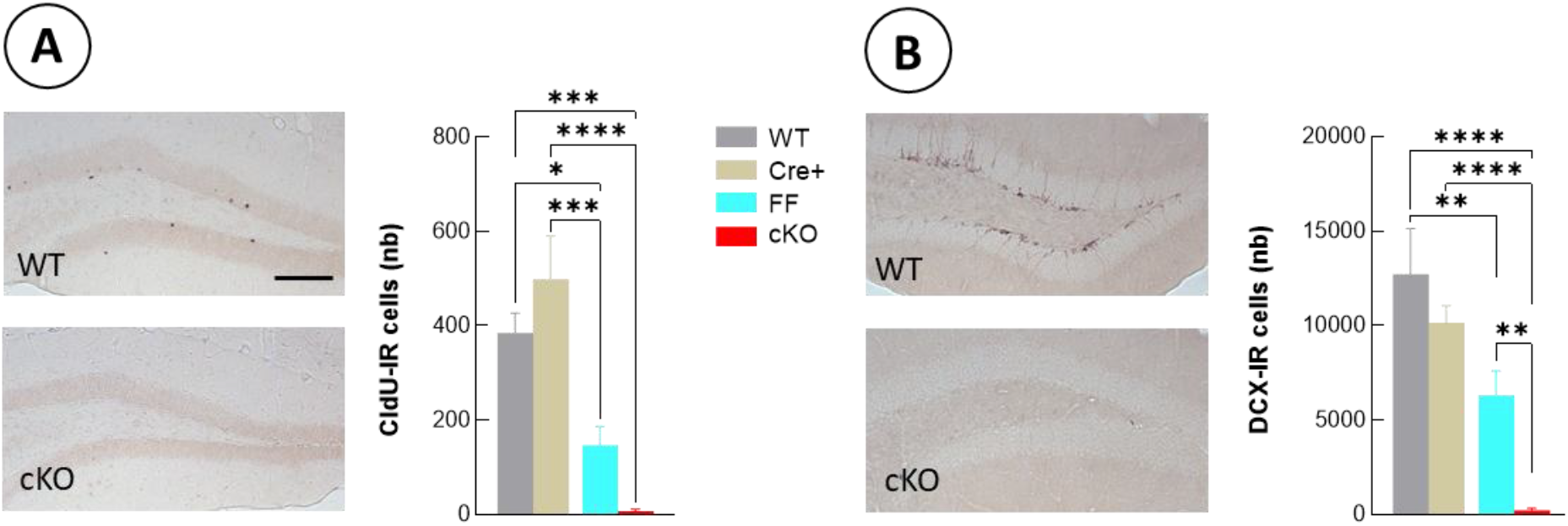
Removal of *Ascl1* in stem cells reduces adult neurogenesis. (A) The total number of CldU-IR cells and (B) the total number of DCX-IR cells was quantified in the granule cell layer (GCL) of the dentate gyrus. Data are mean ± SEM. Scale bar = 100 µm.

Next, the impact of removing *Ascl1* in dentate gyrus stem cells was examined using doublecortin (DCX) as a surrogate of immature neurons. In line with previous results (Andersen *et al*., 2014;Castro *et al*., 2011), the number of DCX-IR cells was significantly lower in FF compared to WT mice (Figure 1B, [Cre effect F_1,24_=9.208, p<0.01; Flox effect F_1,24_=32.5, p<0.0001, Cre x Flox interaction F_1,24_=1.517, p=ns with cKO different from WT, Cre+, and FF mice at p<0.01]).

To verify that the drastic reduction in the generation of adult-born cells did not impact the activity of developmentally-born dentate neurons, we exposed animals to a novel environment (Erwin *et al*., 2020;Sun *et al*., 2021) and neural activity was evaluated using cFos as a proxy. We found as expected a higher level of activation in animals exposed to a novel environment compared to animal remaining in their homecage (Figure S1), and no impact of the genotype in either condition [Homecage, Cre effect F_1,32_=0.2685, p=0.607; Flox effect F_1,32_=3.300, p=0.078, Cre x Flox interaction F_1,32_=0.1547, p=0.696; Novel environment Cre effect F_1,35_=0.3525, p=0.556; Flox effect F_1,35_=1.198, p=0.281, Cre x Flox interaction F_1,35_=2.093, p=0.1569]. Altogether this indicates that any behavioral consequence of removing *Ascl1* in stem cells is likely not sustained by an indirect modification in the activity of developmentally-born cells.

Next, in order to test the consequences of removing adult-born neurons in reconstructing a complete memory from partial cues, we measured pattern completion abilities two months after Tam injection in the Morris water maze (Figure 2A). After a familiarization phase, mice were trained to find a hidden platform from different departure points. During this phase (sessions 1 to 4, Figure S2), all mice, whatever their genotype, learned to find the hidden platform (session effect F_3,99_=15.504; p<0.01; Performance during the 4 sessions: Cre effect F(_3,99_) = 1.024; p = 0.386, Flox effect F_3,99_ = 0.156; p = 0.926, Cre x Flox interaction F(_3,99_) = 0.568; p = 0.637).

**Figure 2.**
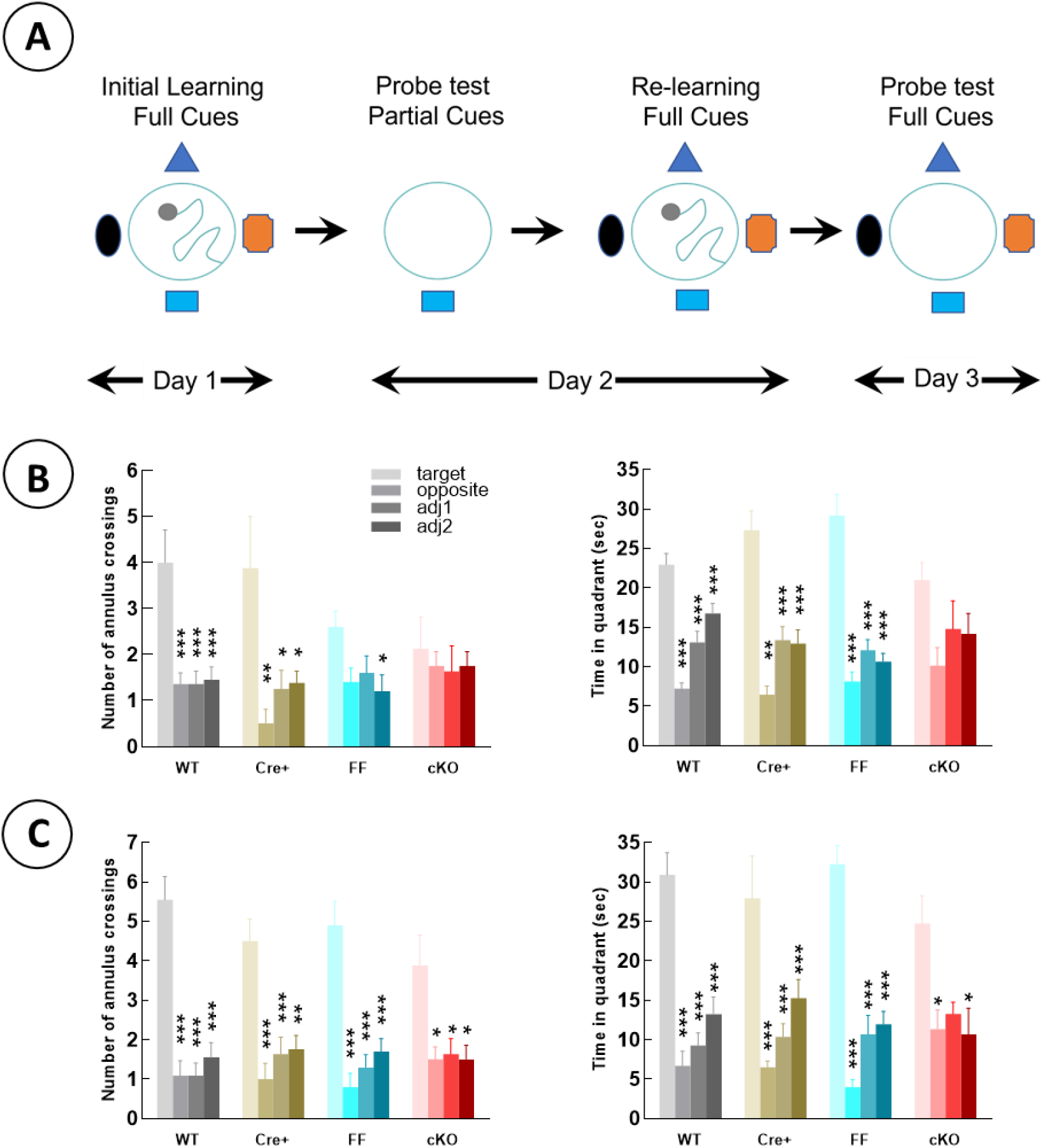
Removal of *Ascl1* in stem cell impairs pattern completion. (A) Experimental protocol. (B) Pattern completion measured during a probe test with partial spatial cues. (C) Spatial probe test when all cues were reintroduced. *** p<0.001, ** p<0.01, * p<0.05 Target vs other.

Then we investigated the role of the adult-born neurons in retrieval in pattern completion condition where three of the principal cues were removed and a probe test was conducted. Analysis of annuli crossings revealed that mice from the four groups did not explore the four quadrants similarly [F_3,99_=14.369; p<0.001]. On the difference in exploration of these 4 quadrants, ANOVA reveals a significant effect of Flox transgene [F_3,99_ = 4.333; p = 0.006] but no effect of Cre transgene [F_3,99_ = 0.302; p = 0.824] and no interaction between Cre and Flox transgenes [F_3,99_ = 0.352; p = 0.788]. A complementary analysis was thus conducted within each group of mice (Figure 2B left panel). A significant quadrant effect was observed in the Control groups [WT: F_3,40_=9.37, p<0.001; Cre+: F_3,28_=4.901, p=0.07] and post-hoc analyses revealed that the target annulus was crossed more frequently than the other three in both groups. In the FF group, a quadrant effect was also found [F_3,36_=3.252, p=0.033], but post-hoc analysis revealed a significant difference only between the Target and Adjacent2 quadrants. Finally, in the cKO mice, no quadrant effect was detected [F_3,28_=0.188, p=0.903], with all four annuli being crossed at similar frequencies.

We also analyzed the overall time in the different quadrants (Figure 2B, right panel), revealing again that mice from the different genotypes do not explore similarly the different quadrants [F_3,99_=45.288, p<0.001]. On the difference in exploration of these 4 quadrants, ANOVA reveals a significant interaction between Flox and Cre transgenes [F_3,99_ = 4.077; p = 0.016] but no main Cre [F_3,99_ = 0.446; p = 0.720] or Flox effect [F_3,99_ = 0.807; p = 0.493]. Thus, mice from the four groups did not appear to explore the four quadrants in the same way. An analysis per genotype revealed a significant quadrant effect in the two control groups and the FF group [WT: F_3,40_=28.206 p<0.001, Cre+: F_3,28_=23.051, p<0.001; FF: F_3,36_=31.622, p<0.001]. Post-hoc analyses revealed that mice spent significantly more time in the Target quadrant than in the other three (p<0.001). In contrast, in the cKO group, there was no quadrant effect [F_3,28_=2.729, p=0.063], revealing a random search in these mice.

Then in a final step, we verified that memory retrieval was preserved when all cues were present. To do so, mice were first re-trained with all spatial cues, followed by a probe test with all the cues. During the relearning phase (Figure S1, sessions 5 to 6), there was no significant change in performance [F_1,33_ = 2.048; p = 0.162]. Analyses showed no significant impact of the genotype of mice [Cre effect F_3,99_ = 1.024, p = 0.386; Flox effect F_3,99_ = 0.156, p = 0.926; Cre x Flox interaction F_3,99_ = 0.568, p = 0.637].

During the probe test with all cues, as expected, there was a significant quadrant effect when anulus crossings or total time in quadrants were considered [Figure 2C left panel, F_3,99_ = 50.391, p < 0.001; Figure 2C right panel, F_3,99_ = 43.404, p < 0.001, respectively for annulus crossings and time in quadrant]. On the difference in exploration of the annuli, we observed no effect of Cre [F_3,99_ = 0.018, p = 0.11], no effect of Flox [F_3,99_ = 0.565, p = 0.639], and no Cre x Flox interaction [F_3,99_ = 0.313, p = 0.816]. For time spent in the different quadrants, a marginal Cre effect was detected [F_3,99_ = 2.728, p = 0.048], but no Flox [F_3,99_ = 0.749, p = 0.525], or Cre x

Flox interaction [F_3,99_ = 1.401, p = 0.247]. A complementary analysis was conducted within each group of mice. A significant quadrant effect was observed in all groups and post-hoc analyses revealed that the mice of the four groups crossed more frequently the target annulus (Figure 2C left panel) and spent significantly more time in the Target quadrant compared to the other three quadrants (Figure 2C, right panel), indicating a good spatial memory.

In the same mice we verified that deficits in pattern completion were not related to motor or motivation deficits by analysis swim speed that was similar between groups during the probe tests. We measured no effect of Cre [F_1,33_ = 0.271, p = 0.606], no effect of Flox [F_1,33_ = 0.329, p = 0.570], and no Cre x Flox interaction [F_1,33_ = 0.707, p = 0.406] in swimming speed.

Finally, in a separate experiment, we verified that memory deficits were not related to an alteration of emotional status (Figure S3; table S1). The impact of removing *Ascl1* in RGLs on anxiety-related behavior was evaluated by measuring avoidance responses to potentially threatening situations. We did not evidence any impact of reducing adult neurogenesis on any of the parameters examined. Concerning depression-related behaviors, anhedonia and resignation were preserved whereas self-grooming behavior was slightly reduced in Flox mice (FF, cKO).

## Discussion

Using an established transgenic mouse model to block the production of adult born neurons, we report for the first time that ablating adult-born dentate granule neurons leads to a specific alteration of pattern completion abilities. Indeed, using both the number of annuli crossing and the time spent in the target quadrant, cKO mice in which adult neurogenesis is almost totally abolished were impaired in reconstructing a complete memory from partial cues in the Morris water maze. These deficits were not due to the speed of pattern completion recall or to anxiety-like or depression-like behavior. Interestingly, a partial reduction of neurogenesis in FF mice was insufficient to disrupt pattern completion, although performances were not as good as controls (annuli crossing).

Previous research has shown that the CA3 region of the hippocampus — the main target of dentate granule cell projections via their mossy fibers (MF) — plays a key role in spatial pattern completion. More specifically, the importance of NMDA receptors in this region has been demonstrated through lesion, genetic, and pharmacological studies (Fellini *et al*., 2009;Lee and Kesner, 2002;Nakazawa *et al*., 2002). In one earlier study, the respective contribution of immature adult-born dentate granule neurons (3–4 weeks old) and mature, or “old,” neurons (>6 weeks old, regrouping developmentally-born and adult-born cells) in pattern completion abilities was already investigated. Authors selectively inhibited MF transmission from most developmentally-born and old adult-born neurons, while leaving the activity of immature ones intact. Because pattern completion recall was disrupted under these conditions, it was concluded that old dentate granule neurons are essential for the recall phase of pattern completion (though not for its overall capacity) (Nakashiba *et al*., 2012).

In the present work, we extend these findings by showing the critical role of adult-born neurons in pattern completion. The observed deficits occurred following a sustained, large-scale reduction of adult neurogenesis over a three-month period. Interestingly, although we found a clear impairment in pattern completion, the speed of recall was unaffected — unlike in the previous study. This difference suggests that developmentally-born neurons may also contribute to pattern completion, possibly in a complementary way, a hypothesis that remains to be experimentally tested.

Our findings have significant implications for aging, during which pattern completion declines. Additionally, maladaptive memories—driven by excessive pattern completion—are central to post-traumatic stress disorder (PTSD) and addiction. In PTSD, minor, non-threatening cues can trigger vivid re-experiencing of traumatic memories, while in addiction, exposure to drug-related cues can induce powerful cravings by reactivating drug-use memories. Given the role of adult-born neurons in these conditions, our discovery suggests that they could constitute potential therapeutic targets to modulate pattern completion, offering new avenues for improving mental health.

## Resource availability

All data are available in the main text or supplementary materials or upon request to the corresponding author.

## Acknowledgments

We thank the animal facility and the genotyping platform of INSERM U1215 NeuroCentre Magendie, funded by INSERM and Labex Brain, for animal care and mouse genotyping. More specifically the help of Cédric Dupuy, Fiona Corailler, and Delphine Gonzales is acknowledged. This work was supported by Institut National de la Santé et de la Recherche Médicale, INSERM (to DNA), Centre National de la Recherche Scientifique, CNRS (to MK), and the Francis Crick Institute, which receives its funding from Cancer Research UK (FC CC2033), the UK Medical Research Council (FC CC2033), and the Wellcome Trust (FC CC2033).

## Author contributions

PR, MK and DNA conceived experiments. PR and PD performed experiments. PR and MK analyzed data. PR, MK and DNA wrote original draft. FG provided biological resources. All authors contributed to the article and approved the submitted version.

## MATERIAL AND METHODS

### A. Animals

Adult male double transgenic mice containing a conditional mutant allele of *Ascl1* and the *Glast-CreERT2* allele were used in the experiments. Ascl1 is a proneural transcription factor, expressed by adult stem cells, promoting their activation, proliferation and differentiation into neurons via the generation of intermediate progenitors (Andersen *et al*., 2014). The conditional mutant allele of *Ascl1* (*Ascl1* ^*fl/fl*^) consists of the coding sequence of *Ascl1* flanked by loxP sites on both ends allowing its deletion by the Cre recombinase. The Cre recombinase is fused with modified estrogen receptor (ERT2) that restricts the enzyme in the cytoplasm, activating its translocation to the nucleus to mediate recombination only upon Tam stimulation (Pacary *et al*., 2011). Cre-ERT2 is expressed under the GLAST promoter, which is a marker for adult neural stem cells. The double transgenic mice containing both the alleles was obtained from the breeding of heterozygous male mice carrying the Glast-CreERT2 construct (Glast-CreERT2 mice) with female mice homozygous for the floxed *Ascl1* construct (*Ascl1* ^*fl/fl*^ mice = FF mice), as previously described (Andersen *et al*., 2014;Pacary *et al*., 2011). Genotypes generated by this breeding and used in the study were *Glast-CreERT2* ^*-/-*^ ; *Ascl1* ^*+/+*^ (WT), *Glast-CreERT2* ^*+/-*^; *Ascl1* ^*+/+*^ (Cre+), *Glast-CreERT2* ^*-/-*^ ; *Ascl1* ^*fl/fl*^ (FF), and *Glast-CreERT2* ^*+/-*^ ; *Ascl1* ^*fl/fl*^ (cKO). All experimental procedures were carried out following guidelines of the European Union (2010/63/UE), and were approved by the ethical committee of Bordeaux (CEEA50; Dir 1399, A16502).

### B. Tamoxifen treatment

Two-month-old mice were administered intraperitoneally (i.p.) with tamoxifen diluted in corn oil (Tam) for 5 consecutive days at a concentration of 100mg/kg body weight.

### C. Injection of CldU

One month after Tam injection, a batch of 28 mice (WT n=7, Cre^+^ n=8, FF n=6, cKO n=7) was injected with Choro-2’-Desoxyuridine (CldU, 42.75 mg/kg dissolved in 0.9% NaCl), once per day during 2 days. Animals were sacrificed three months after Tam injection, i.e. 2 months after CldU injection.

### D. Baseline and novelty-induced activity of granule cells

In this experiment, 75 mice were treated with Tam at 2-months of age. Two months later they were housed in individual cages, and 6 months later, they were sacrificed under baseline conditions (HomeCage: WT n=10, Cre^+^ n=7, FF n=10, and cKO n=9) or 2 hours after exposure to a new bedding-free cage (Novel Environment: WT n=11, Cre^+^ n=7, FF n=11, and cKO n=10).

### E. Immunohistochemistry and stereology

Animals were anesthetized and perfused transcardially with 0.1 M phosphate buffered saline (PBS, pH 7.4), followed by 4% buffered paraformaldehyde (PFA). Brains were collected and post-fixed in PFA at 4 °C for a week. Subsequently, 40 μm-thick coronal sections were cut using a vibratome (Leica) and stored in cryoprotectant medium (30% ethylene glycol, 30% glycerol in KPBS) at −20 °C before staining.

Free-floating sections were processed in a standard immunohistochemical procedure in order to visualize CldU (Anti-BrdU, 1/1000, Accurate OBT0030), doublecortin (DCX, 1:8000; Sigma D9818), or cFos (cFos 1/1000; Cell Signaling Technology 2250S)-labeled cells. Briefly, sections for CldU staining were exposed to HCl (2N) for 30 min at 37°C and rinsed with PBS, while for DCX and cFos, sections were directly rinsed in PBS. Then all sections were treated with methanol and 0.5% H2O2 for 30 min. Sections were washed again in PBS before incubation with a blocking solution containing 3% normal serum and 0.3% Triton X100 in PBS for 45 min at room temperature. They were then incubated for 48 h at 4 °C with the primary antibodies diluted in the blocking buffer. Then sections were incubated with biotin-labeled secondary antibodies diluted in PBS-1% normal serum, and immunoreactivities were visualized by the biotin–streptavidin-peroxydase technique (ABC HRP kit; Dako) with 3,3′-diaminobenzidine (DAB) as chromogen. The number of immunoreactive (IR) cells throughout the entire granule and subgranular layers of the left DG was estimated using the optical fractionator method as previously described(Koehl *et al*., 2021).

### F. Pattern completion

#### 1. Animals

In this experiment, 37 mice were treated with Tam at 2-months of age (WT n=11, Cre^+^ n=8, FF n=10, and cKO n=8). Two months later, they were trained in the water maze.

#### 2. Task apparatus

The water maze was an ivory circular pool (110 cm diameter, 30 cm high) filled to a depth of 15 cm with water maintained at 23 ± 1°C. The water was made opaque by addition of a white opacifier. A white-painted platform (9 cm diameter) was placed inside the pool, 15.5 cm away from the pool wall. The apparatus was surrounded by white curtains preventing mice from using visual extra-maze cues and was illuminated by three lamps heterogeneously placed behind the curtains(Florian and Roullet, 2004). Four white and black visual landmarks approximately located at 50 to 100 cm from the pool, differing in shapes, motives and size, were placed on the curtains (Fig. 1A). The swimming pool was surmounted by a video camera connected to a video recorder and a computerized tracking system (Ethovision®, Noldus).

#### 3. Behavioral Procedures

On day 0, mice were individually submitted to a single familiarization session of three trials with the platform located always in the centre of the pool and protruding 0.5 cm over the surface of the water. The starting positions were determined in a pseudorandom order, such that each of them was used once in a single session. If the mouse did not find the platform within one minute, it was gently guided to it. The animal stayed one minute on the platform between the trials and before returning to the home cage. All mice showed similar latencies in finding the visible platform [data not shown, F(2,34) = 0.232; p = 0.794].

On day 1, mice were given four consecutive acquisition sessions of three trials with an inter-session delay of 25-30 min during which they were returned to their home cage. The procedure was the same as in the familiarization phase, except that the platform was placed in a specific quadrant (target quadrant) and was submerged 0.5 cm beneath the surface of the water.

On day 2, the platform was removed from the pool and mice were submitted to a 60-sec probe-trial. During this first probe trial (PT1), the three largest patterns were removed, thus only the smallest one was maintained. Thirty min after this probe trial, mice received 2 sessions of reacquisition in the standard condition with the platform in the target quadrant and with all patterns around the pool.

On day 3, mice were submitted a second 60-sec probe-trial (PT2), but this time, all cues were present.

### G. Emotional behaviors

#### 1. Animals

In this experiment, 48 mice were treated with Tam at 2-months of age (WT n=13, cre+ n=10, FF n =11, and cKO n=14). Behavioral analysis began 2 months after Tam injection. Mice were placed into individual cages one week before the first test session.

#### 2. Anxiety-related behaviors

The impact of removing *Ascl1* in RGLs was first examined on anxiety-related behavior by measuring avoidance responses to potentially threatening situations, such as unfamiliar open environments(Koehl *et al*., 2021). Mice were first tested in an *open-field* (OF; 50 x 50 cm white PVC square arena; ∼400 lux) to evaluate locomotor performance and exploratory activity. Mice were placed in the center of the OF and allowed to explore the test for 10 min. Total distance traveled and time spent in the corners (the safest place of the open-field) were recorded and analyzed using a videotracking system (© Videotrack, ViewPoint).

Two days later, a *light/dark emergence test* was conducted in the same open-field (∼400 lux) containing a cylinder (10 cm deep, 6.5 cm diameter in dark gray PVC) located length-wise along one wall, its opening facing a corner of the OF. Mice were placed into the cylinder and tested for 15 min. Total time spent inside the cylinder and total number of entries in the cylinder (safe place) were analyzed.

Finally, an *elevated plus maze* (EPM) test was performed 48h later. The test was a transparent Plexiglass apparatus with 2 open (45 x 5 cm) and 2 closed (45 x 5 x 17 cm) arms extending from a common central platform (5 x 5 cm). Mice were released from the central zone, facing an open arm and allowed to freely explore the maze for 5 min (∼50 lux). Time spent in and total number of entries into the open and closed arms were measured. Standard measures of rodent anxiety were calculated: % time and % entry in the open arms compared to time in open arms + closed arms, and total entries into any arm of the maze; in addition, total distance traveled in the open and closed arms was taken as a measure of activity/exploratory tendency.

#### 3. Depression-related behaviors

Depression-related behaviors were measured in the same animals upon completion of the anxiety tests as previously described (Koehl *et al*., 2021). Briefly depression-related behaviors were examined by measuring avolition (lack of motivation or inability to initiate goal-directed behavior) in the nest building and sucrose splash tests, anhedonia in the sucrose preference test, and resignation/behavioral despair in the Forced swim test (FST). At least 4 days separated each test.

***Nest building*** was measured by placing a cotton nestlet in each cage in the morning and scoring nest quality 24 h later.

*The* ***sucrose splash test*** was carried out by spraying a high viscosity 10% sucrose solution on the coat of the mice to induce a self-grooming behavior. Latency to initiate the first grooming episode, as well as frequency of grooming over a 5-min period was measured immediately after applying the solution.

For the ***sucrose preference*** test, mice were first habituated for 48 h to the presence of two drinking bottles filled with tap water. They were then given, for 48 h, a free choice between one bottle filled with a 4% sucrose solution, and the other with tap water. Sucrose intake was calculated as the amount of consumed sucrose in mg per gram body weight, and sucrose preference was calculated according to the formula: sucrose preference = (sucrose intake)/(sucrose intake + water intake) × 100.

The ***Forced swim test (FST)*** was performed last in a 5L glass cylinder filled with 26±1°C water to a depth of 20 cm so that mice cannot touch the bottom of the cylinder. One 6-min session was run and both latency to float and duration of immobility over the last 4 min of the test were scored by an experimenter unaware of the experimental groups.

### H. Statistical analysis

Statistical analyses were performed with GraphPad Prism 10 software (GraphPad v10.0.3; GraphPad Software LLC). No statistical methods were used to predetermine sample sizes. Investigators were blind to group allocation in behavioral experiments. Graphs represent mean values ± SEM. A 95% confidence interval was used for statistical comparisons.

---

**Figure S1:**
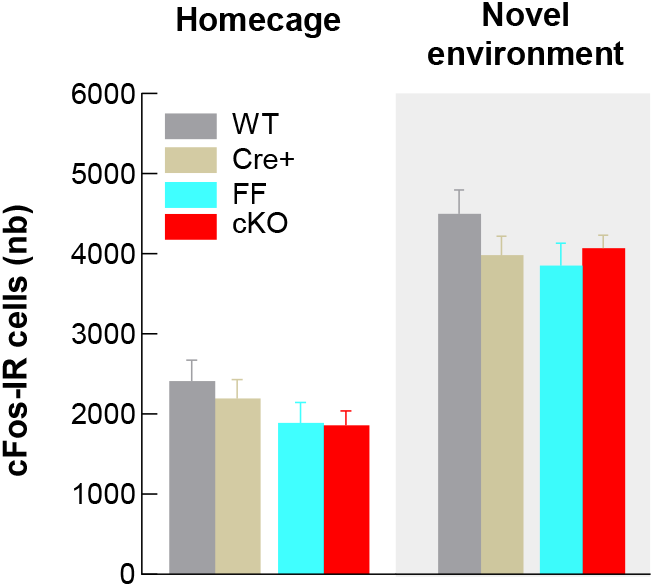
Removal of *Ascl1* in stem cell does not modify baseline or novelty-induced cFos expression in DG.

**Figure S2:**
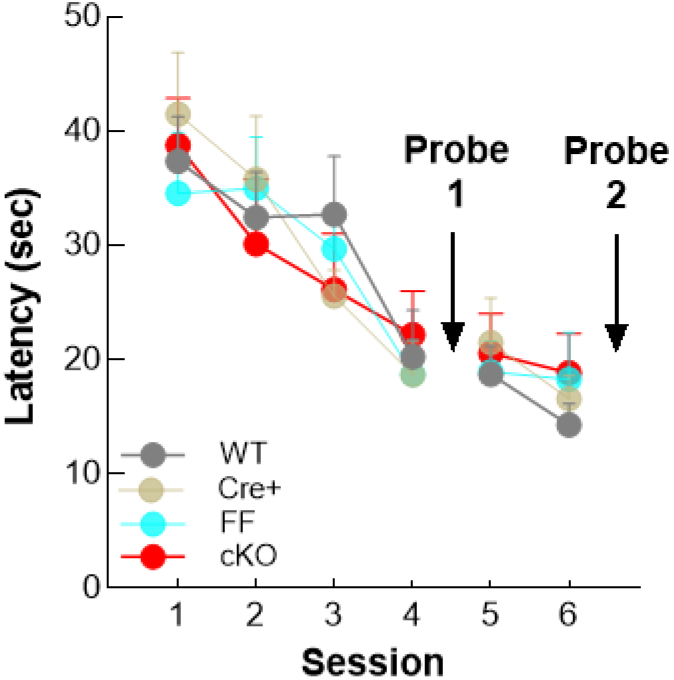
Initial learning and re-learning with all spatial cues. Probe 1: first probe trial with partial cues. Probe 2: second probe trial with partial full cues.

**Figure S3.**
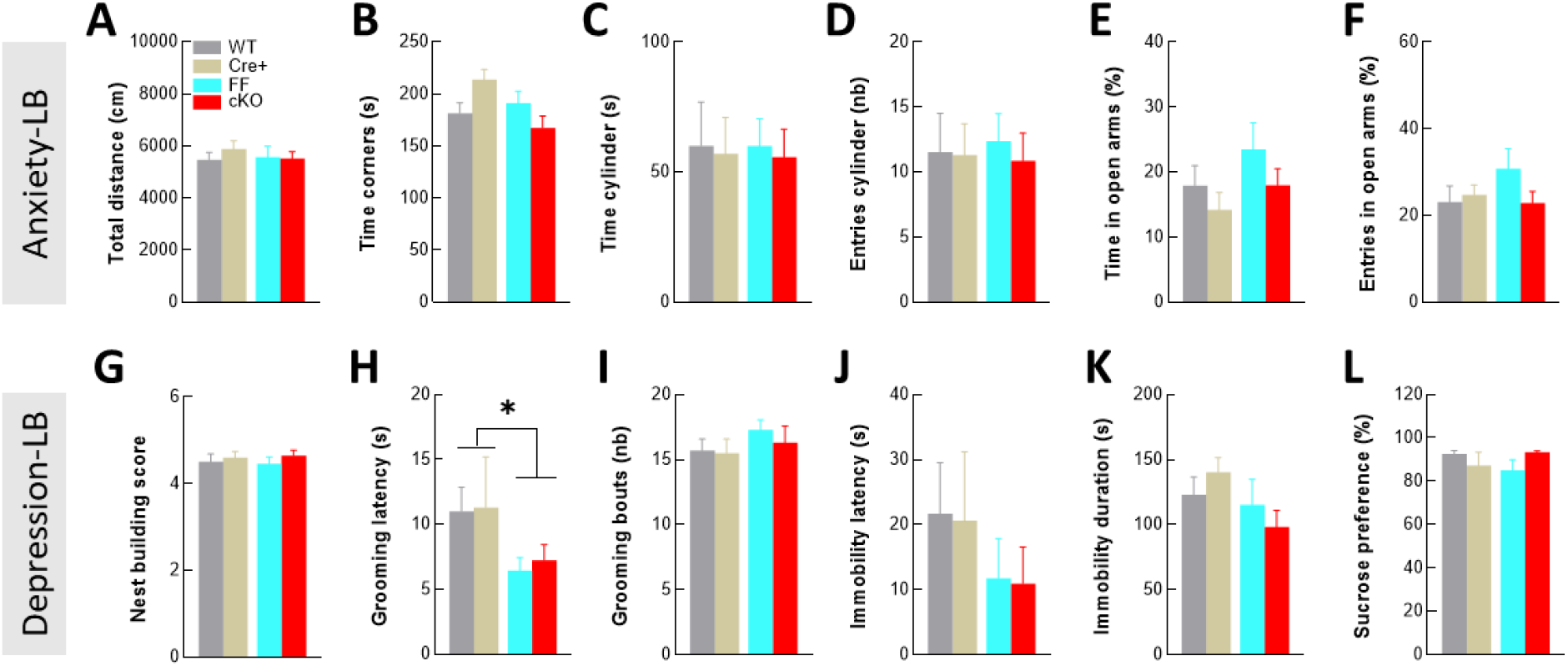
Removal of *Ascl1* in stem cells does not influence anxiety and depression-like behavioral responses. Anxiety-like responses were measured in the open-field (A**-B**), in the light/dark emergence task (C, D) and in the elevated plus-maze (E, F). Depression-like responses included measures of *motivation* evaluated in the nest building (G) and the sucrose splash tests (H, I), of *resignation* evaluated in the forced swim test (J, K), and of *anhedonia* measured in the sucrose preference test (L).

**Table S1:** Statistical analysis of anxiety and depression-like behavioral responses.

| Test | Variable | Anova effect | F(DFn, DFd) | P value |
| --- | --- | --- | --- | --- |
| Open-Field | Total distance<br>(Fig S3A) | Cre | F (1, 44) = 0,3084 | P=0,5815 |
|  |  | Lox | F (1, 44) = 0,1737 | P=0,6789 |
|  |  | Interaction | F (1, 44) = 0,5129 | P=0,4777 |
|  | Time corners<br>(Fig S3B) | Cre | F (1, 44) = 0,1504 | P=0,7000 |
|  |  | Lox | F (1, 44) = 2,726 | P=0,1058 |
|  |  | Interaction | F (1, 44) = 6,491 | P=0,0144 |
| Light-dark emergence test | Time Cylinder<br>(Fig S3C) | Cre | F (1, 44) = 0,07217 | P=0,7895 |
|  |  | Lox | F (1, 44) = 0,002263 | P=0,9623 |
|  |  | Interaction | F (1, 44) = 0,001756 | P=0,9668 |
|  | Entries cylinder<br>(Fig S3D) | Cre | F (1, 44) = 0,1216 | P=0,7290 |
|  |  | Lox | F (1, 44) = 0,005837 | P=0,9394 |
|  |  | Interaction | F (1, 44) = 0,06421 | P=0,8011 |
| Elevated plus maze | Time open arms<br>(Fig S3E) | Cre | F (1, 44) = 2,170 | P=0,1478 |
|  |  | Lox | F (1, 44) = 2,208 | P=0,1444 |
|  |  | Interaction | F (1, 44) = 0,08437 | P=0,7728 |
|  | Entries open arms<br>(Fig S3F) | Cre | F (1, 44) = 0,8138 | P=0,3719 |
|  |  | Lox | F (1, 44) = 0,6848 | P=0,4124 |
|  |  | Interaction | F (1, 44) = 1,899 | P=0,1751 |
| Nest building | Nest score<br>(Fig S3G) | Cre | F (1, 44) = 0,8395 | P=0,3645 |
|  |  | Lox | F (1, 44) = 6,814e-005 | P=0,9935 |
|  |  | Interaction | F (1, 44) = 0,07877 | P=0,7803 |
| Sucrose splash test | Grooming latency<br>(Fig S3H) | Cre | F (1, 44) = 0,06407 | P=0,8013 |
|  |  | Lox | F (1, 44) = 4,250 | P=0,0452 |
|  |  | Interaction | F (1, 44) = 0,01206 | P=0,9131 |
|  | Grooming bouts<br>(Fig S3I) | Cre | F (1, 44) = 0,3035 | P=0,5845 |
|  |  | Lox | F (1, 44) = 1,222 | P=0,2750 |
|  |  | Interaction | F (1, 44) = 0,1378 | P=0,7122 |
| Forced swim test | Immobility latency<br>(Fig S3J) | Cre | F (1, 44) = 0,01438 | P=0,9051 |
|  |  | Lox | F (1, 44) = 1,672 | P=0,2027 |
|  |  | Interaction | F (1, 44) = 0,0002052 | P=0,9886 |
|  | Immobility duration<br>(Fig S3K) | Cre | F (1, 44) = 1,261e-005 | P=0,9972 |
|  |  | Lox | F (1, 44) = 2,899 | P=0,0957 |
|  |  | Interaction | F (1, 44) = 1,395 | P=0,2439 |
| Sucrose preference test | Sucrose preference<br>(Fig S3L) | Cre | F (1, 44) = 0,1811 | P=0,6725 |
|  |  | Lox | F (1, 44) = 0,03955 | P=0,8433 |
|  |  | Interaction | F (1, 44) = 3,751 | P=0,0592 |

